# Regional brain development analysis through registration using anisotropic similarity, a constrained affine transformation

**DOI:** 10.1101/574129

**Authors:** Antoine Legouhy, Olivier Commowick, Maïa Proisy, François Rousseau, Christian Barillot

**Affiliations:** Univ Rennes, CNRS, INRIA, INSERM, IRISA UMR 6074, Empenn ERL U-1228, F-35000, Rennes, France; CHU Rennes, Radiology Department, F-35033 Rennes, France; IMT Atlantique, LaTIM U1101 INSERM, UBL, Brest, France

## Abstract

We propose a novel method to quantify brain growth in 3 arbitrary orthogonal directions of the brain or its sub-regions through linear registration. This is achieved by introducing a 9 degrees of freedom (dof) transformation called anisotropic similarity which is an affine transformation with constrained scaling directions along arbitrarily chosen orthogonal vectors. This gives the opportunity to extract scaling factors describing brain growth along those directions by registering a database of subjects onto a common reference. This information about directional growth brings insights that are not usually available in longitudinal volumetry analysis. The interest of this method is illustrated by studying the anisotropic regional and global brain development of 308 healthy subjects betwen 0 and 19 years old. A gender comparison of those scaling factors is also performed for 4 classes of age. We demonstrate through these applications the stability of the method to the chosen reference and its ability to highlight growth differences accros regions and gender.

## Introduction

In pediatric image analysis, the study of brain development provides insights in the normal trend of brain evolution and enables early detection of abnormalities. Many types of morphometric measurements based on structural images have been explored and have shown their reliability as biomarkers in clinical use as established in [1]. Evaluated on a database of subjects covering a period of interest, it allows to better model the brain development and to highlight changes in growth, shape, structure, etc. Those measurements can be conducted on geometrical objects of different dimensions. Unidimensional ones such as the bicaudate ratio (minimum intercaudate distance divided by brain width along the same line) have been explored in [2] and [3] but also biparietal, bifrontal and transverse cerebellar diameters in [4], as well as more areal quantities such as cortical surface in [5] or corpus callosum mid-sagittal area in [6]. However, the vast majority of studies are based on 3D features through the assessment of region of interest (ROI) volumes. Volumetry measures of different regions of the brain have been considered for specific ages or various temporal ranges. A far from exhaustive list is presented in Figure 1. Studied regions are very heterogeneous from large areas such as the whole brain itself, cerebellum, lobes or partitions of those to smaller ones such as basal ganglia, hippocampus, thalamus sometimes even separated according to the composition of their tissues (white matter (WM), gray matter (GM), cerebro-spinal fluid (CSF)). Some group comparisons have also been performed mostly between male and female or between preterm and term newborns.

**Fig 1.**
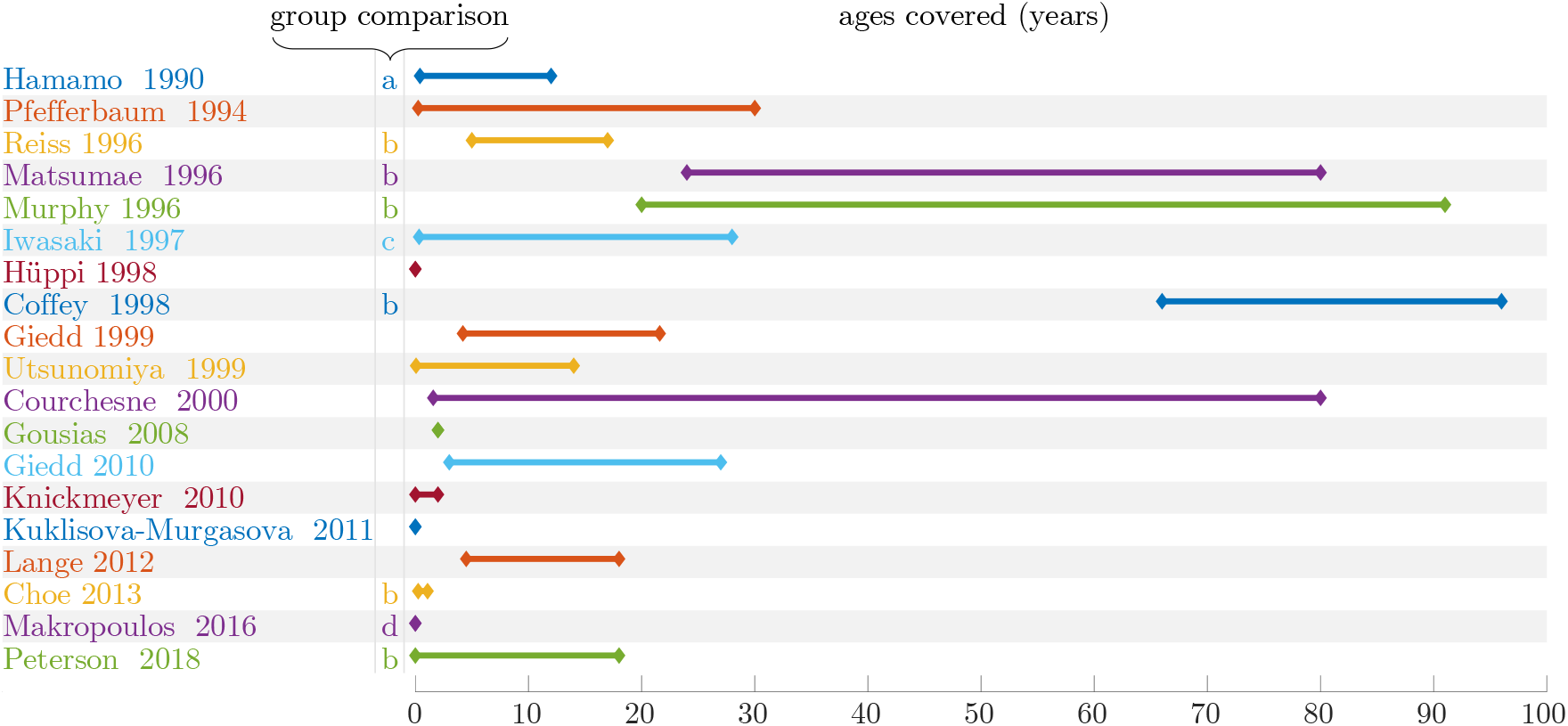
Ages covered by different brain volumetry sudies, group comparison between: a - normal / mental retardation, b - male / female, c - normal / motor disturbances, d - normal / premature. [7], [8], [9], [10], [11], [12], [13], [14], [15], [16], [17], [18], [6], [19], [20], [21], [22], [23], [24]

Morphometric measurements can be determined manually. However, this requires the intervention of a medical expert able to select specific landmarks or perform segmentation. These tasks are highly time consuming with a potentially large inter-expert variability. Advances in computational medical imaging allow nowadays the use of semi-automated (requiring some human intervention) or fully-automated techniques. They involve algorithms able to automatically perform operations such as registration and segmentation.

A major drawback of purely volumetric measurements is that they do not provide any information on the shape of the regions or about the anisotropy of their development. In this paper, a new method is proposed that aims at quantifying global and regional brain growth in three arbitrary orthogonal directions of the brain (or ROI) through linear registration. To do so, a transformation called anisotropic similarity is introduced in section 1. It is an affine transformation with scaling directions constrained by orthogonal vectors arbitrarily chosen. A method to estimate, in a 3 dimensional space, the optimal anisotropic similarity for the least squares problem of distances between two sets of paired points is presented in section 2. Those results will then be used to create a registration algorithm based on this transformation. By registering a database of subjects onto a common basis (i.e. an atlas segmented in different ROIs) using anisotropic similarity, we have the opportunity to extract global or regional scaling ratios for all those subjects along arbitrary chosen orthogonal directions.

A direct application is, using the pipeline exposed in section 3, the exploration of regional scaling ratios growth charts along three fixed orthogonal directions through the ages highlighting anisotropic brain development. Resulting curves for whole brain and ROIs (lobes, basal ganglias, cerebellum…) are presented in section 4.2. A comparison of scaling factors from males and females is performed for 4 different classes of age between 0 and 19 years old in section 4.3. Finally, the influence of the common reference image on the resulting scaling factors is studied in section 4.4.

Anisotropic similarity registration algorithm as well as other image processing tools used in this paper are publicly available in Anima^1^ (open source software for medical image processing).

## 1 Theoretical background

### 1.1 Generalities about linear transformations and anisotropic similarity especially

An affine transformation is a composition of a linear map *A* (*N* × *N* matrix) and a translation *t* (in ℝ^*N*^) operating on coordinates: *y* = *Ax* + *t*. Using singular value decomposition (SVD) on *A*, we obtain:

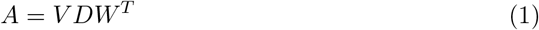

where *W* and *V* are unitary matrices and *D* is a positive diagonal matrix. By introducing *R* = *V*. Det(*V*), *U* = *W*. Det(*W*) and *S* = Det(*V*) Det(*W*)*D*, we get a modified decomposition:

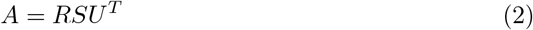

where *U* is a rotation matrix defining the directions of scaling, *S* is a diagonal scaling matrix and *R* is a rotation matrix. We define a new linear transformation, hereafter named anisotropic similarity, which is an affine transformation with constrained directions of scaling. In other words, we define an anisotropic similarity transformation as an affine one where *U* is fixed. Summing up, we have the following in 3D space:

- An affine transformation has 12 degrees of freedom:

– a rotation (3 dof): the matrix *U* determines scaling directions.
– an anisotropic scaling (3 dof): matrix *S*.
– a rotation (3 dof): matrix *R*.
– a translation (3 dof): vector *t*.
- For an anisotropic similarity, the directions of scaling defined by *U* are constrained. This leaves 9 dof: 3 for rotation, 3 for scaling and 3 for translation.
- For a similarity, the scaling part is constrained to have identical values on the diagonal i.e. *S* = *s* Id with *s* ∈ ℝ leading to a linear part of the form *sRU^T^*. This leaves 7 dof: 3 for rotation, 1 for scaling and 3 for translation.
- For a rigid transformation, the scaling part is constrained to identity leading to a linear part of the form *RU^T^* which is a rotation matrix since rotation matrices form a group for matrix multiplication. This leaves 6 dof: 3 for rotation and 3 for translation.

### 1.2 Generalities about linear registration

Registration consists in finding an optimal transformation that matches a moving image onto a reference image. This transformation is usually obtained by maximizing a similarity criterion. Many rigid (or linear in general) registration methods have been developed. They can be divided into two families: the ones that try to match geometrical features such as contours or surfaces, and those called iconic that are based on voxel intensities. Some of them use a global similarity measure between the two images such as mutual information in [25] and [26], while others rely on local similarities. Among this second category of approaches, block matching strategies exposed in [27] and [28] have gained in popularity. In those methods, two steps are iterated:

1. Matching: for a set of blocks established in the reference image, homologous blocks best satisfying a similarity criterion are searched in the moving image.
2. Aggregation into a global transformation: an optimization is performed in order to find the global transformation minimizing a distance between the sets of blocks and is then applied to the moving image. Usually, the weighted sum of squared euclidean distance is chosen for the cost function.

In order to perform an anisotropic similarity registration using the block-matching method, the two steps mentioned above have to be iterated. The first one (matching) is performed the same way it would be for any regular linear transformation. It outputs two sets of paired points: *x* and *y* that are in our case the centers of the homologous blocks. The second step (aggregation onto a global transformation) however is dependent on the type of linear transformation we want to determine leading to an adapted optimization in each case.

This optimization step consists in finding, in the set of eligible transformations, the one that best maps *x* onto *y*. Let *x* = {*x*_1_,…, *x_M_*} and *y* = {*y*_1_,…, *y_M_*} be two sets of *M* paired points coming from the matching step. For a global transformation with linear part *A* and translational part *t*, the least squares problem associated to the matching of *x* and *y* consists in the minimization of the following criterion:

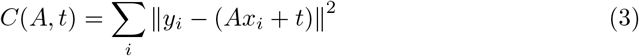

#### Remark.

For the sake of clarity we presented a version with a non-weighted least squares problem but the reasoning is the same with a weighted one.

The optimal translation 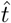 can be directly obtained from the optimal linear part (independently of the type of linear transformation) from the barycenters of the two sets of points as developed in [29]. Let 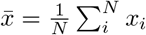 and 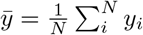, we have then:

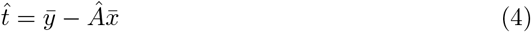

Let 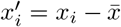 and 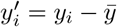 be the barycentric coordinates, the problem can then be simplified as:

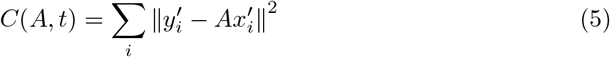

In the case of the linear part being affine, there is no constraint. A closed form solution can therefore be easily found as shown in [29]. For rigid and similarity transformations, constraints lead to more complicated lagrangians but a closed form solution can be found as well using unit quaternions in 3D space as a representation of rotations like in [30] and [29].

## 2 Optimal anisotropic similarity between two sets of paired points

To our knowledge, the optimization procedure in the case of anisotropic similarities has not been considered in the literature. We thus present a method also based on quaternions to find the optimal anisotropic similarity between two sets of paired points. Writing A as its decomposition, the goal is to minimize the following criterion:

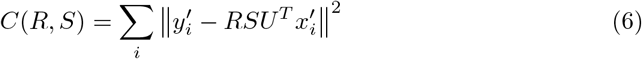

Where *U* is fixed. Let 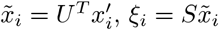

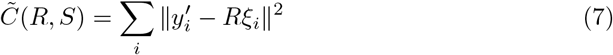

R can be expressed using quaternions following [30] and [29] and the problem then becomes (see A.1):

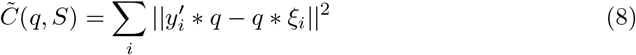

Where *q* is a unit quaternion and * is the quaternion multiplication. Let *p* and *q* be quaternions. There is a matricial representation of quaternions allowing to express quaternion product as a matrix product. Matricial quaternions *P* and *Q* are defined such that: *Q_p_q* = *p* * *q* and 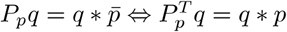.

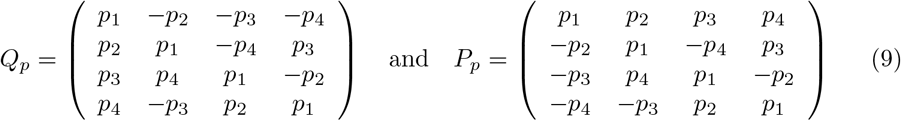

Using those matricial quaternions on 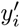 and ξ*_i_* taken as pure quaternions, we have 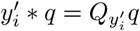 and 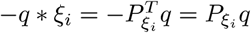. Thus, we obtain the following criterion (see A.2):

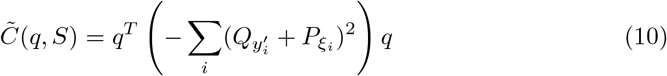

For further computation, we denote 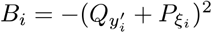 and 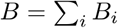. A lagrangian with unit constraint *q^T^q* = 1 has then to be added to ensure a unit quaternion:

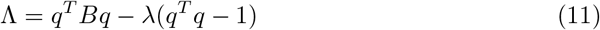

The derivatives of this new formulation can then be written as:

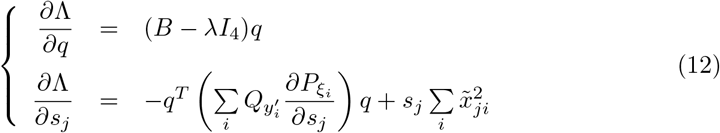

Derivative with respect to *q* depends upon *s_j_* and vice versa. Therefore, a direct solution to the problem of minimizing 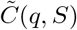 is difficult to find if not impossible. However, separating the problem between *S* and *q* leads to an alternate optimization scheme, each having an analytical solution.

*Rotation*:

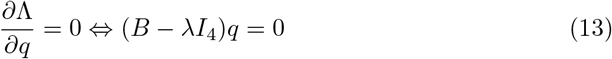

Solving this equation amounts finding the eigen vectors of *B*. More precisely, the global minimum 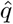 is the one associated to the smallest eigen value of *B* as shown in [30], [29]. *Scaling*: (see A.3)

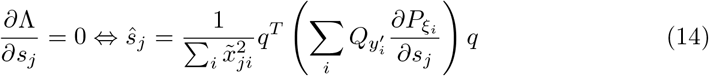

Now, interestingly the matrices 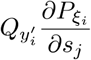 have a quite trivial form. They are all symmetric, only the placing and indexes change (see A. 4). We finally get the followin iterative alternate optimization scheme:

- For a fixed value of *Ŝ*, estimate the new optimal rotation quaternion: 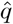 as the eigenvector with the smallest eigenvalue of *B*
- For a fixed value of 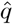, estimate the new optimal scaling matrix *Ŝ* = Diag(*ŝ*_1_, *ŝ*_2_, *ŝ*_3_) following:

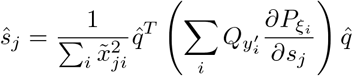

## 3 Material and methods

In the previous section, a method to find the optimal anisotropic similarity between two sets of paired points has been depicted. This gives the opportunity to register a database of subjects onto a common reference image using this type of linear transformation to extract scaling factors along chosen orthogonal directions and to study the variation of these factors on different ROIs between populations or among normal subjects.

### 3.1 Material

308 T1-weighted images of healthy subjects between 0 and 19 years old have been used, coming from three different studies: ASLpedia (section 6.1.1), C-MIND (section 6.1.2) and the Developing Human Connectome Project (dHCP) (section 6.1.3). Details on age repartition among databases and on image characteristics are given in Figure 2.

**Fig 2.**
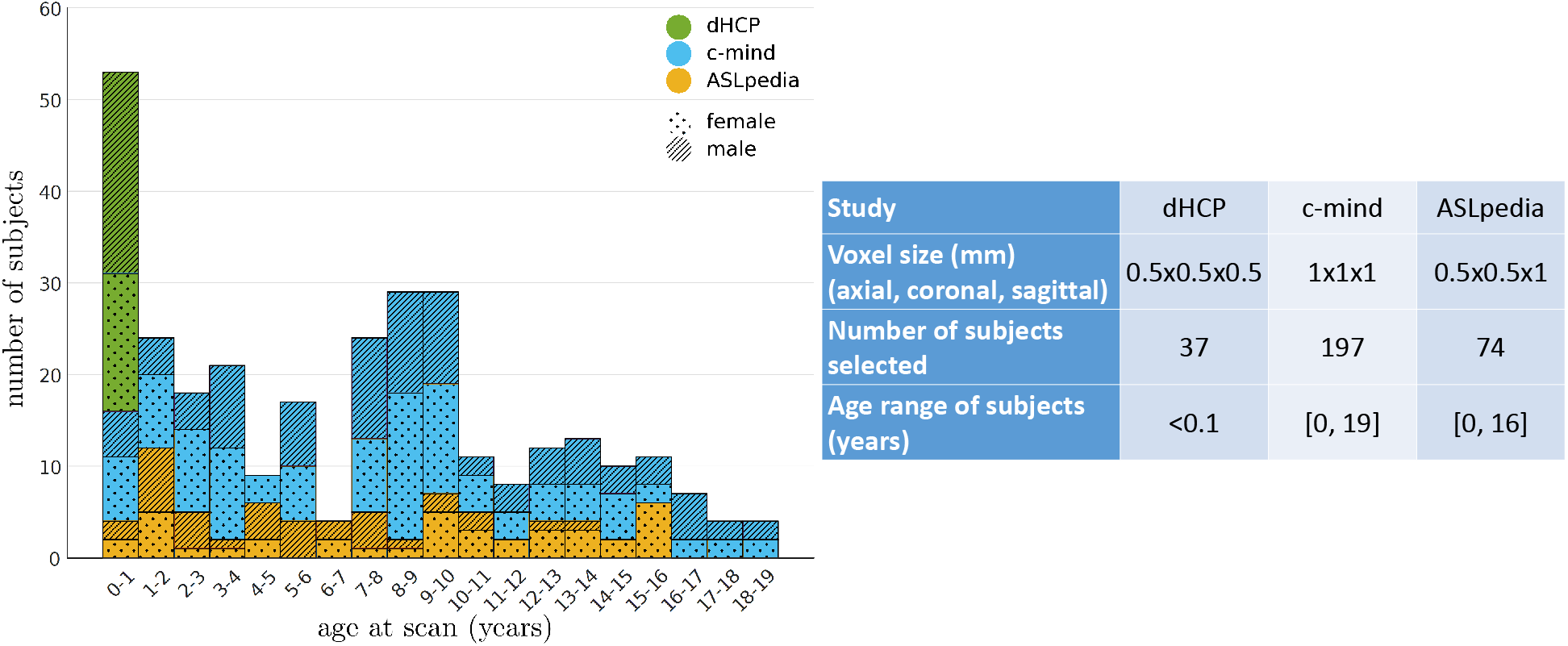
Repartition of the subjects selected from three studies over age

### 3.2 Methods

We developed a pipeline composed of 5 steps to extract scaling factors for 3 orthogonal directions on ROIs from a database of subjects.

1. Choice and construction of the common reference image
2. Segmentation of the common reference image into different ROIs
3. Choice of the constrained directions of scaling for the anisotropic similarity registration
4. Anisotropic similarity registration of a database of subjects to each ROIs of the common reference image to extract relative scaling factors
5. Renormalization of the relative scaling factors to obtain absolute scaling factors

The above numbers associated to the steps are also associated to the subsections numbering below and to the numbers in Figure 3.

**Fig 3.**
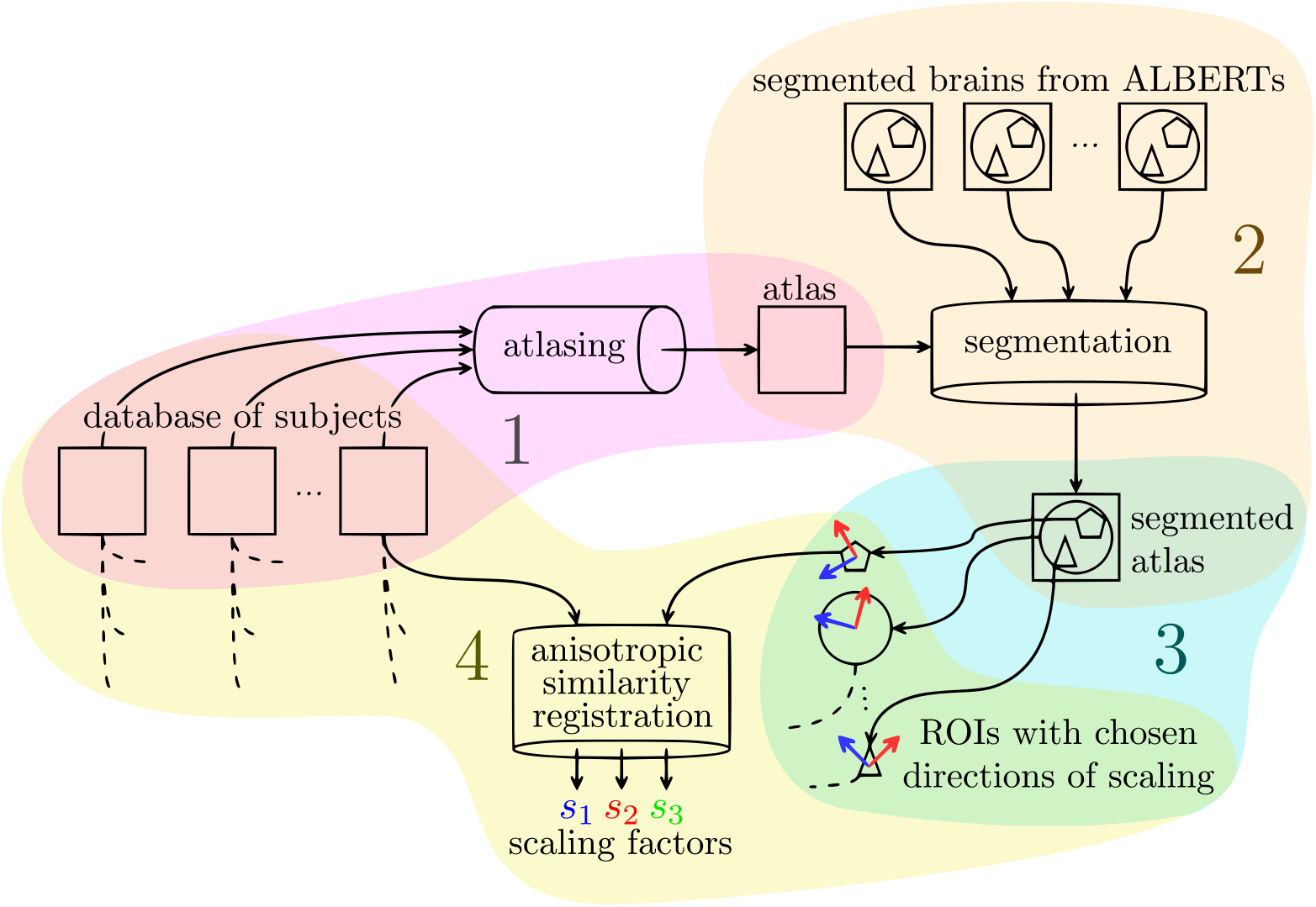
Pipeline for the extraction of scaling factors of a database of subjects using anisotropic similarity registration onto an atlas based on them as common reference image

#### 3.2.1 Creation of the common reference image

For genericity, the common reference image has been chosen to be an atlas made from all the subjects using a modified version of the atlas creation algorithm from [31] available in Anima-Scripts^2^ (open source scripts for medical image processing). The original method computes an atlas up to an affine transformation, biased in that sense by the reference image. This is due to the fact that, in the process, only the residual local transformations are averaged, ignoring global affine ones. Our method, developed in [32], takes advantage of the log-Euclidean framework developed in [33] and the Baker-Campbell-Hausdorff formula, mentionned in [34] and [35], allowing to average the composition of an affine transformations and a diffeomorphism. This adjustment leads to the creation of atlases up to a rigid transformation.

#### 3.2.2 Segmentation of the common reference image

The atlas has then been segmented into 21 regions of interest (ROIs): whole brain, hemispheres, frontal lobe, parietal lobe, temporal lobe, occipital lobe, basal ganglias, cerebellum, insulas, ventricules, corpus callusum and brainstem. All structures were also separated in their left and right sides. To do this segmentation, ALBERTs manual ones ([36] and [37], see acknowledgments 6.1.4) have been used: 20 brains segmented into 50 regions manually drawn based on MRI brain scans that we fused to obtain the wider desired regions. The T1 weighted images of those brains have been registered onto our atlas through affine then diffeomorphic registration. The outputs have then been used to transfer all the segmentations onto our atlas which have been then merged using majority voting following [38]. The segmented atlas is shown Figure 4.

**Fig 4.**
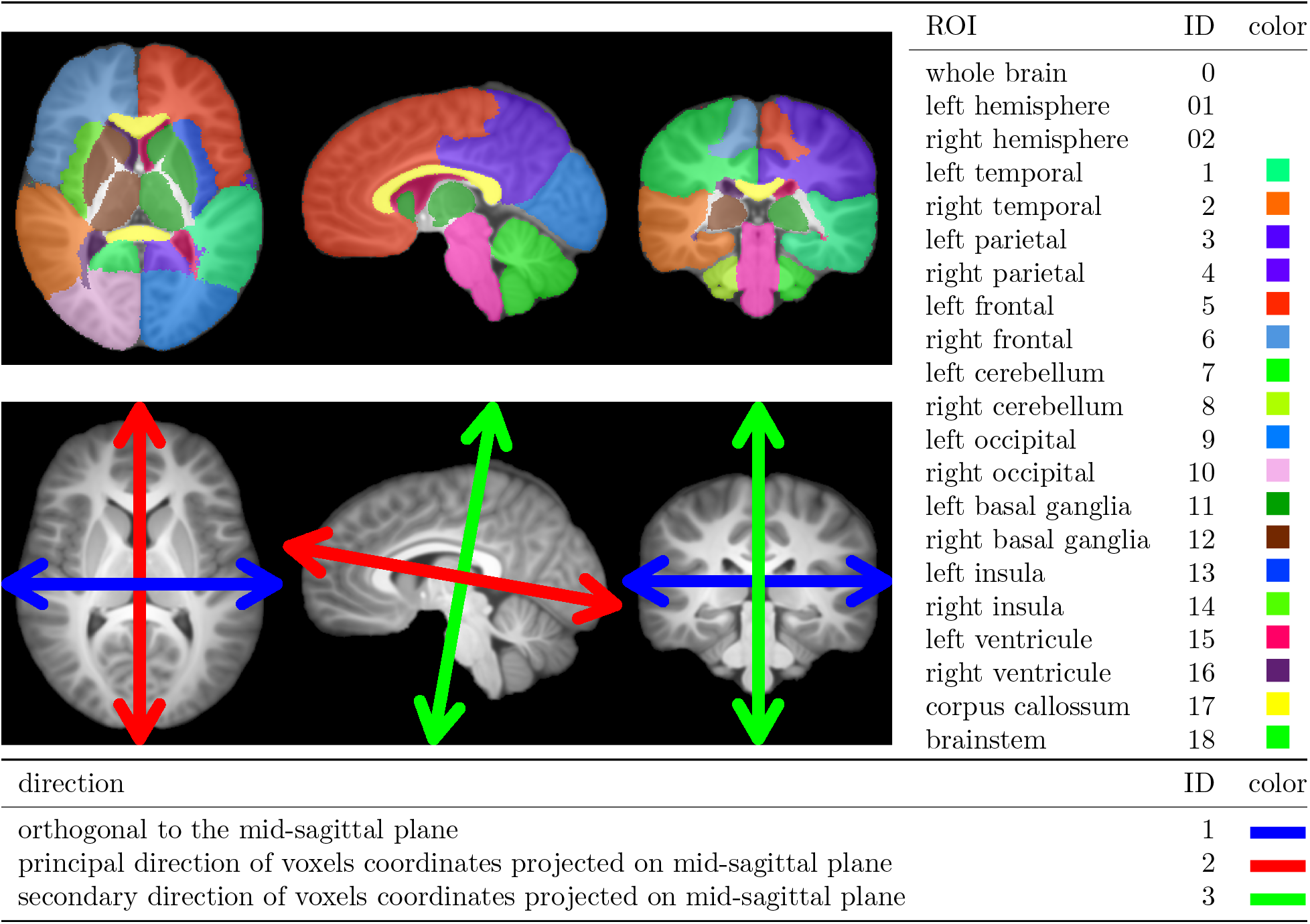
Regions of interest (ROIs) segmented and represented on the common reference image (top), chosen directions of scaling for anisotropic similarity registration defined and represented on the common reference image (bottom)

#### 3.2.3 Choice of the constrained directions of scaling

The fixed scaling directions (characterized by the column vectors of the matrix *U*) are chosen on the reference image such that the first direction (blue in figure 4) is orthogonal to the mid-sagittal plane (determined using [39]) for symmetry reasons. The others two directions are set using principal component analysis (PCA) on the non zero voxels coordinates projected onto the mid-sagittal plane. The second direction (red in figure 4) corresponds to the principal direction from the PCA while the third (green in figure 4) corresponds to the secondary one. 3 orthogonal directions are now chosen: one through iconic considerations and the other ones based on purely geometrical features. In our application, the matrix *U* is the same for all ROIs of the reference image and is defined using the whole brain. However, it is possible to define a different *U* for each ROI independently. Chosen directions of scaling are shown Figure 4.

#### 3.2.4 Anisotropic similarity registration

For each ROI, all subjects undergo an anisotropic similarity registration onto the reference image masked by this ROI. This registration is performed in two steps using in each case our block matching algorithm implemented in Anima^3^:

1. A similarity from whole brain subjects onto whole brain common reference is first estimated.
2. An anisotropic similarity initialized from the previous step output is then computed to bring the subjects onto the atlas masked by the current ROI.

The first transformation, a similarity, is computed indirectly during a process of affine registration. Let *A* be the linear part of an affine transformation *T_A_*. We consider the following SVD: *A* = *VDW^T^* with *D* diagonal positive, *V* and *W* unitary matrices. We define *T_B_* (the nearest similarity associated to *T_A_*) as the transformation with linear part 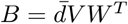 with 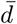 being the average of the singular values namely the mean of the diagonal of *D*, and translation part 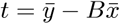. We chose the initialization to be a similarity since the composition of a similarity *T_B_* and an anisotropic similarity *T_C_* associated to a matrix *U* is still an anisotropic similarity associated to the same *U* : *T_B_T_C_* = (*s_B_R_B_*)(*R_C_S_C_U^T^*) = (*R_B_R_C_*)(*s_B_S_C_*)*U^t^* = *RSU^T^*.

**Fig 5.**
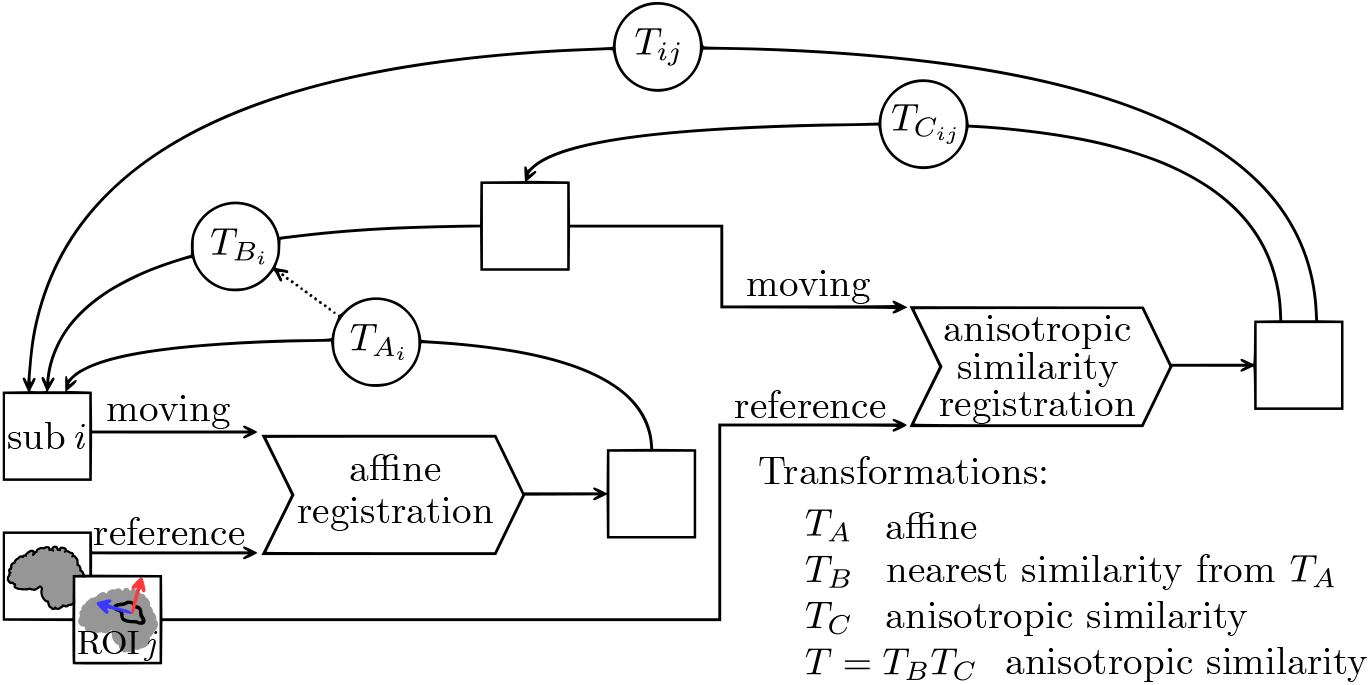
Two steps registration process: first an affine from which a nearest similarity is deduced, then an anisotropic similarity

##### Remark.

Transformations are composed and represented with arrows from destination to start since the interpolations occurring in the resampling process are done using the backward mapping. The inverse transformation is actually used on each voxel of the output images to determine the position in the input image from which a value is sampled.

#### 3.2.5 Extraction of absolute scaling factors

From the output transformations of the registration step, the relative scaling parameters along the three fixed directions are extracted. A normalization is then applied such that the scalings at age 0 are close to 1. To this end, the fact that all dHCP subjects are very young (less than 1 month) is exploited. All relative scaling factors are divided by the average of the ones associated to the dHCP subjects considered as the “root” of the brain expansion. Those new scaling factors will now be considered as absolute scaling factors. At this stage, for each subject, an absolute scaling factor has been determined for each ROI. Those absolute scaling factors are used to model the expansion of the brain toward the chosen directions.

## 4 Experiments and results

### 4.1 Model selection

Several models are traditionally used to represent growth in biostatistics such as the exponential or Weibull models. The second one has been considered by [24] as the best suited to model brain growth in terms of volume. Our case however is different: it can be viewed as a 3-way unidimensional approach. In our quest to find the function best suited to model growth curves for our data, we decided to consider, as a prior, that the brain expansion is stopping at some point. Therefore, we restricted the spectrum to functions that have an horizontal asymptote at infinity. The selected candidates to model brain growth in the chosen directions are the following:

- Rational with polynomials of degree 1 as numerator and denominator : 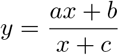
- Weibull: *y* = *a* – *be*^−*cx^d^*^
- Gompertz: *y* = *ae*^−*be*^−*cx*^^
- Exponential: *y* = *a* + *be*^−*cx*^

For each candidate, the optimal coefficients are estimated through nonlinear regression using the Levenberg-Marquardt iterative weighted least squares algorithm from [40]. In this process, weights are chosen to compensate for local gender repartition. For each subject *i*, a window of width *l* = 2 years centered on the subject age is considered. Let *u_f_, n_m_* and *n* be the number of female, male and total subjects respectively in that window. A correction coefficient 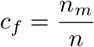 is applied if *i* is a female and 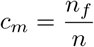 if *i* is a male. Let {*y*_1_,…, *y_n_*} be the observations (i.e. here the obtained scaling factors), 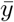 be the average of those and {*ŷ*_1_,…, *ŷ_n_*} be the fitted values.

Based on these statistics, the chosen candidate for the modeling will be the one that best satisfies a criterion quantifying the goodness of fit. This indicator should evaluate the accuracy of the model i.e. how close the model is to the observation while discouraging overfitting. It therefore consists in a tradeoff between accuracy and parsimony. It has been shown in [41] that the coefficient of determination is not, at least when considered alone, an appropriate measure for the goodness of fit in the case of nonlinear model selection. A more adapted goodness of fit for nonlinear model selection is the Akaike information criterion (AIC) developed in [42] and [43]. Based on information theory, it proposes to estimate the information loss induced by each candidate model to represent an unknown process that supposedly generated the data as shown in [44]. This is made possible through the estimation of the Kullback-Leibler divergence related to the maximized log-likelihood. AIC is defined by: 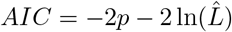, where 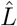 is the maximum likelihood of the candidate model and the first term penalizes a large number of parameters *p*. Therefore, the preferred model among the candidates is the one with the lower AIC. Note that AIC of a model taken alone is meaningless, it makes sense only when compared to the one of the other models. This is why it is recommended to consider it along with an other statistic that quantifies the error between the model and the data like mean of squared errors (MSE): 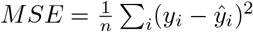 which is the average of the residuals. A corrected version of the AIC has been developed to avoid overfitting in the case of small sample sizes: 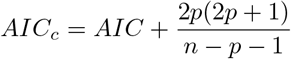. To facilitate the interpretation that can be quite obscure using raw AIC, following [45], it is possible to transform those values into conditional probabilities for each model called Akaike weights. Defined for each model *i* by 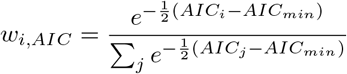, those weights represent the probability for each candidate *i* to be the best suited in the sense of AIC to model the data among all the candidates.

All the goodness of fit depicted above as well as MSE have been evaluated for each of our candidates to model the scaling factors for each ROI. We present the results of this evaluation Figure 6. The Gompertz and exponential models are largely below the other two. Even though the Weibull model behaves relatively well, the rational one shows better scores whatever the tested goodness of fit.

**Fig 6.**
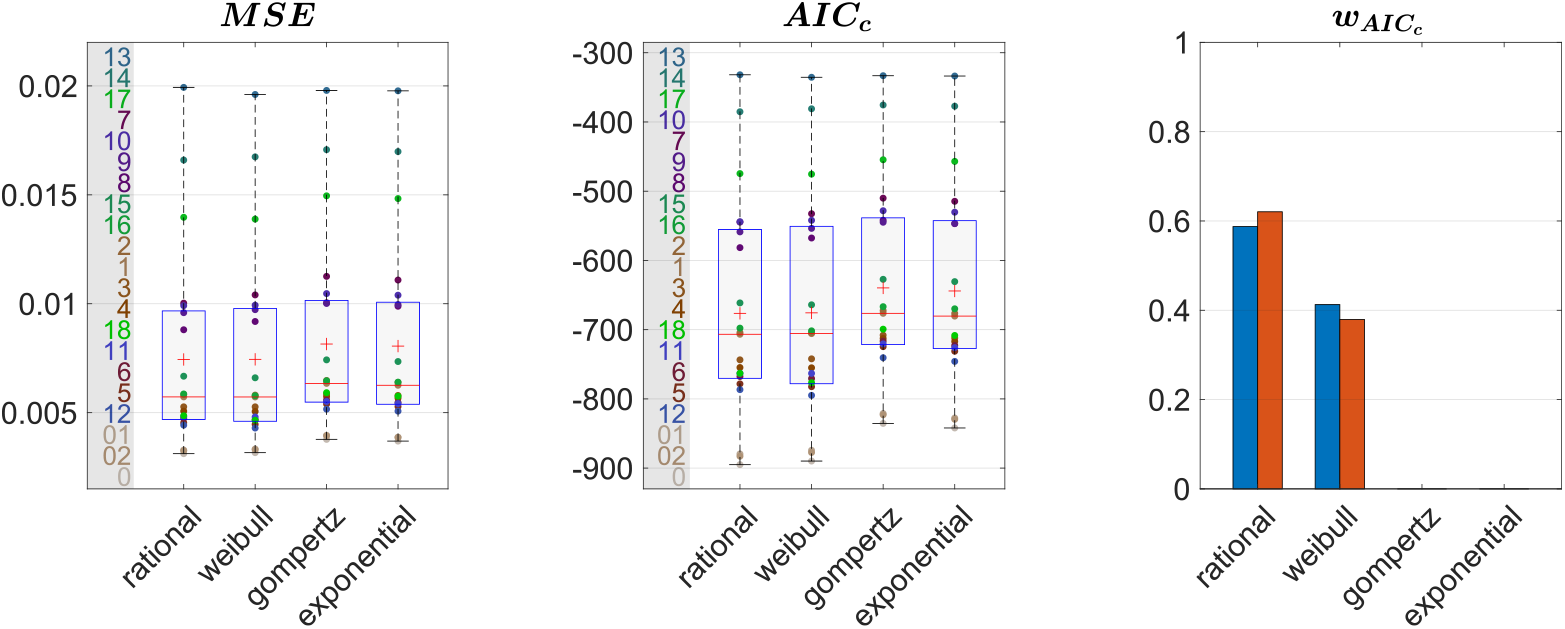
Goodness of fits for each candidate to model the ouputed scaling factors averaged in the 3 directions. Boxplots are performed along the ROIs, ROI IDs are displayed on the left. Akaike weights are computed on mean (blue) and median (red) AIC_c_

### 4.2 Directional growth curves

From the previous sections, scaling factors in each direction for each ROI are now modeled using a rational function with polynomials of degree 1 as numerator and denominator chosen after model selection. Results for all regions studied are presented in figure 7. The method presented by [46] is used to compute simultaneous 99% confidence intervals for fitted values. The black curve represents the average brain growth computed as the mean of the directional models (Figure 7).

**Fig 7.**
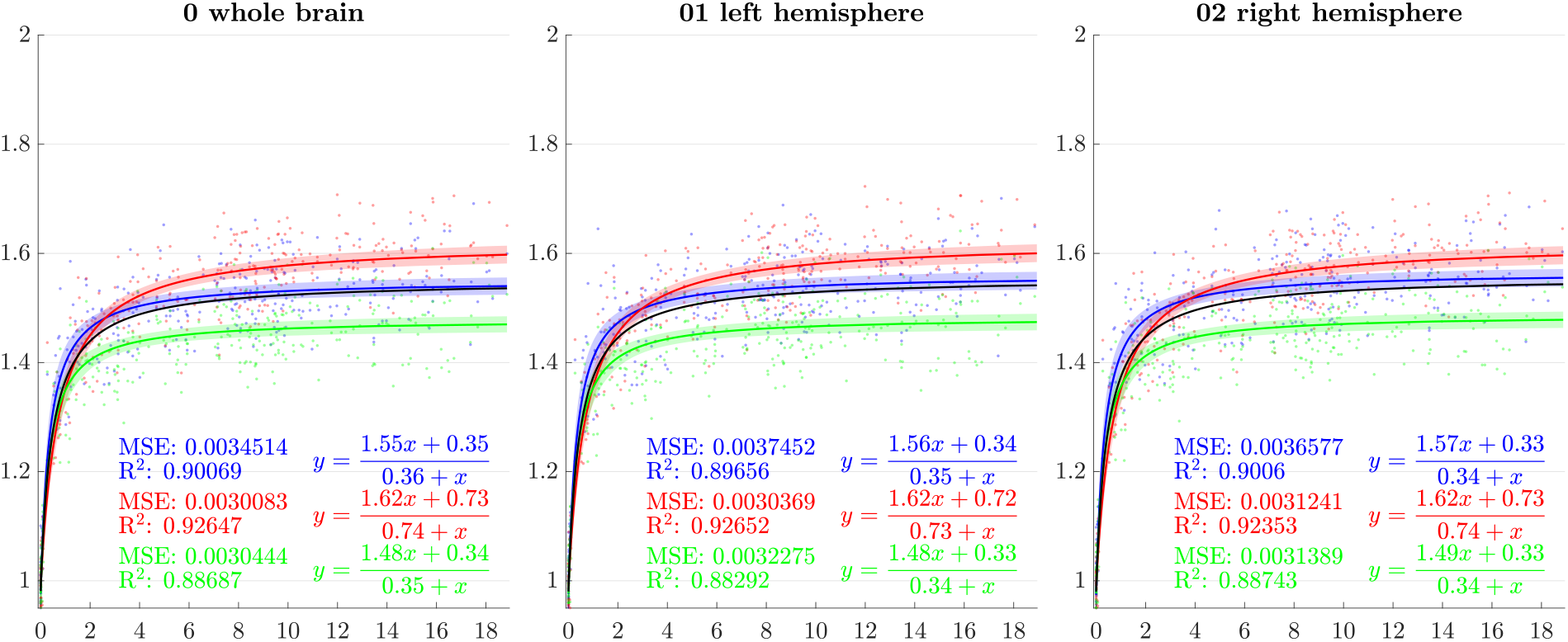
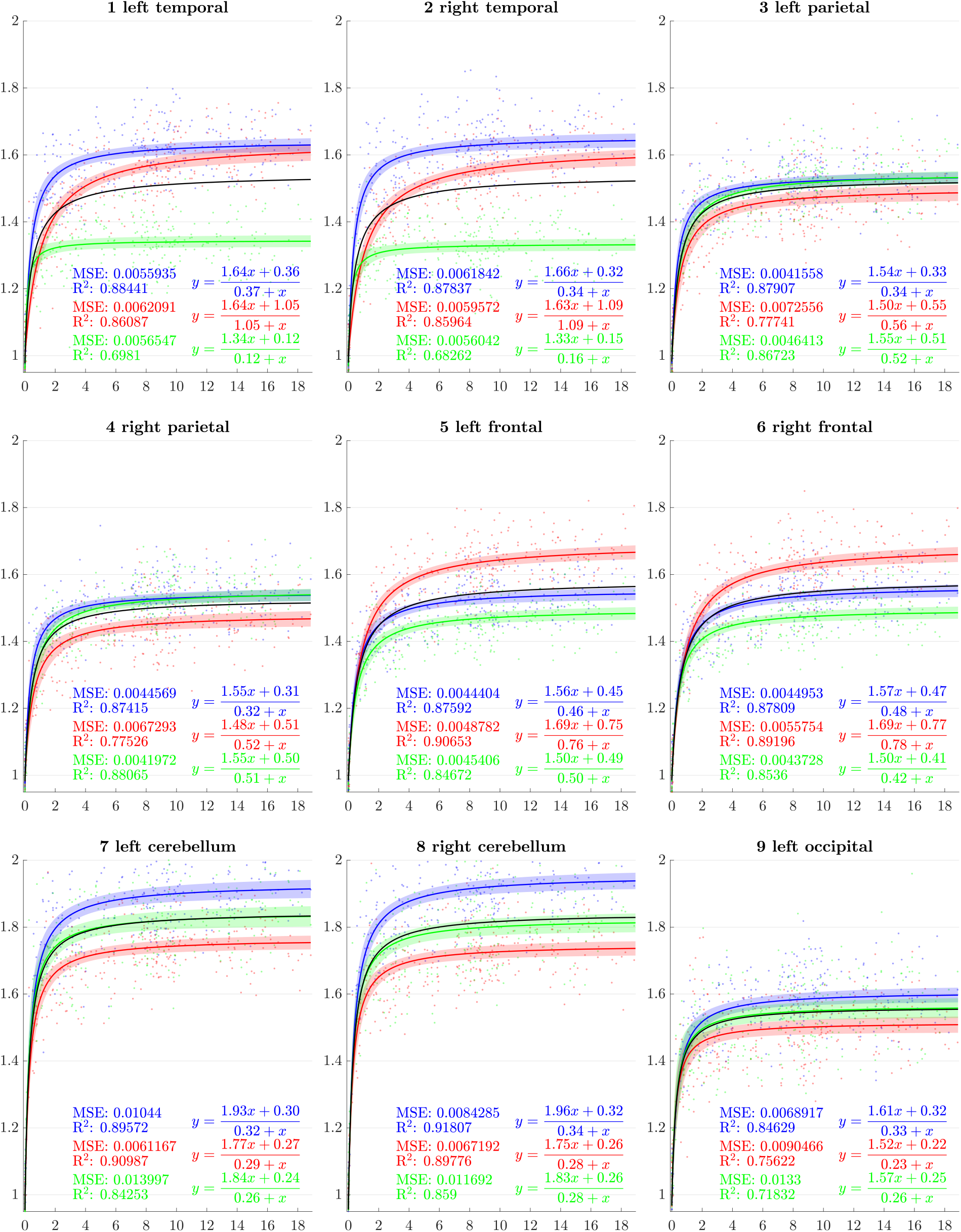
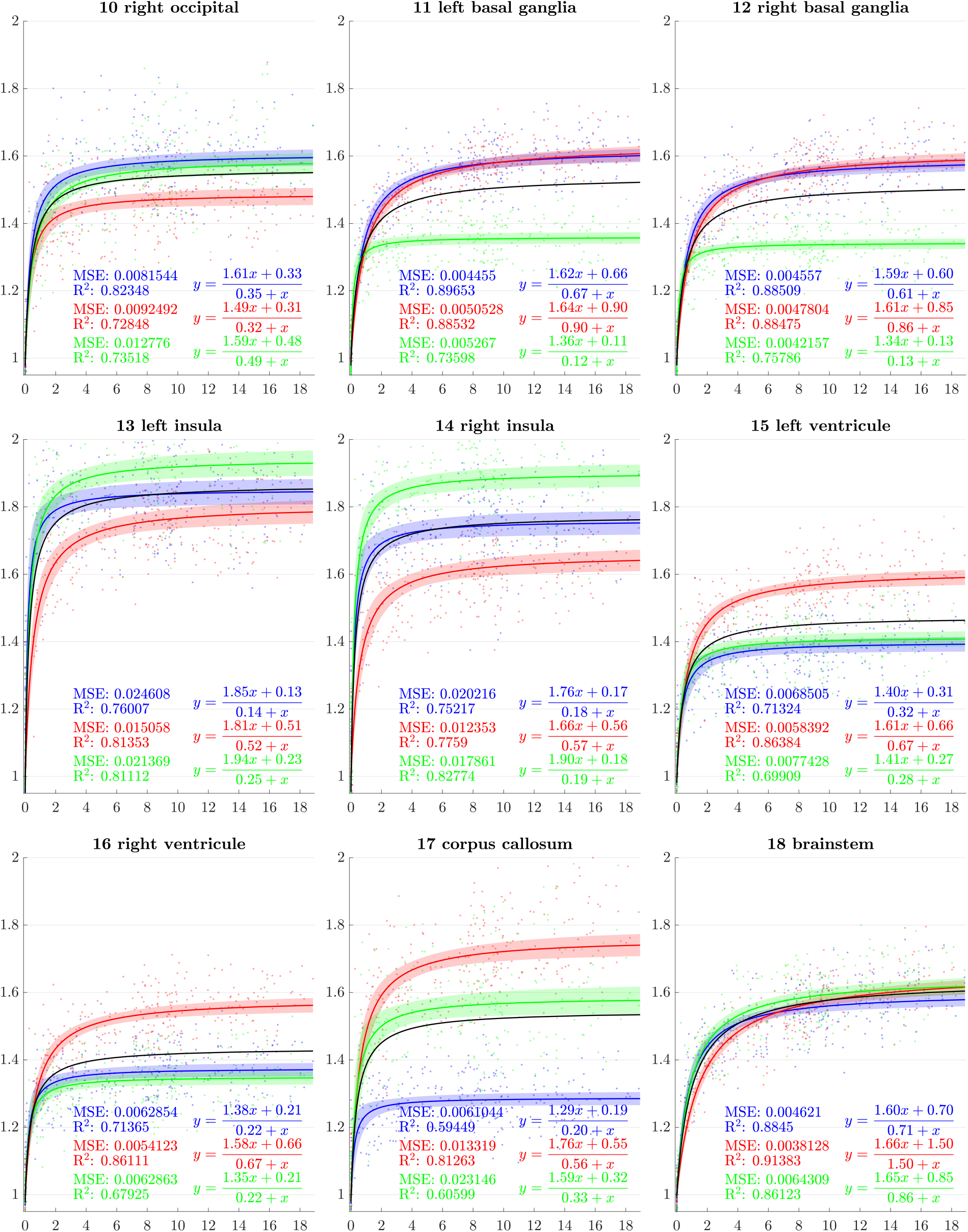
Resulting scaling factors as a function of the age for differents ROIs, along direction 1 (blue), 2 (red), 3 (green). Fitted using rational model together with 99% confidence intervals for fitted values.

### 4.3 Male vs female comparison

Gender, like age, is a characteristic of the subjects available in all the studies we considered. The aim of this section is to evaluate if differences in terms of scaling factor can be found between genders. We divided our data into four classes based on the age of the subjects. The first one contains dHCP participants (newborns), the second one is composed of all non-newborn subjects between 0 and 6 years old, the third one between 6 and 12 and the fourth superior to 12 years old. Repartition of the subjects in terms of gender, age class and study is shown Table 1.

**Table 1.**
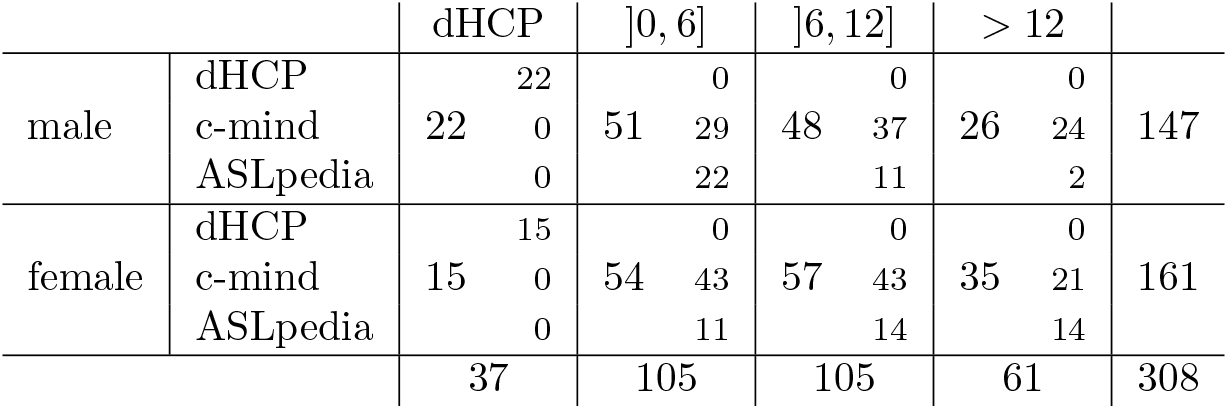
Repartition of the subjects in term of age class, gender and study

For each of these classes, and each of the chosen scaling directions, and each ROI, we performed a test to evaluate if the scaling factors for male subjects are greater than scaling factors for female subjects. Since these data are not normally distributed in those subdivisions, we used two-tailed Wilcoxon-Man-Whitney U-tests. For each of those tests, the null hypothesis *H*_0_ is the following: the distribution of the scaling factors between males and females are equal, while the alternative hypothesis *H*_1_ states: the distributions of males and females are different. We performed 252 tests in total: 4 age classes × 21 ROIs × 3 directions whose results are shown figure 8.

**Fig 8.**
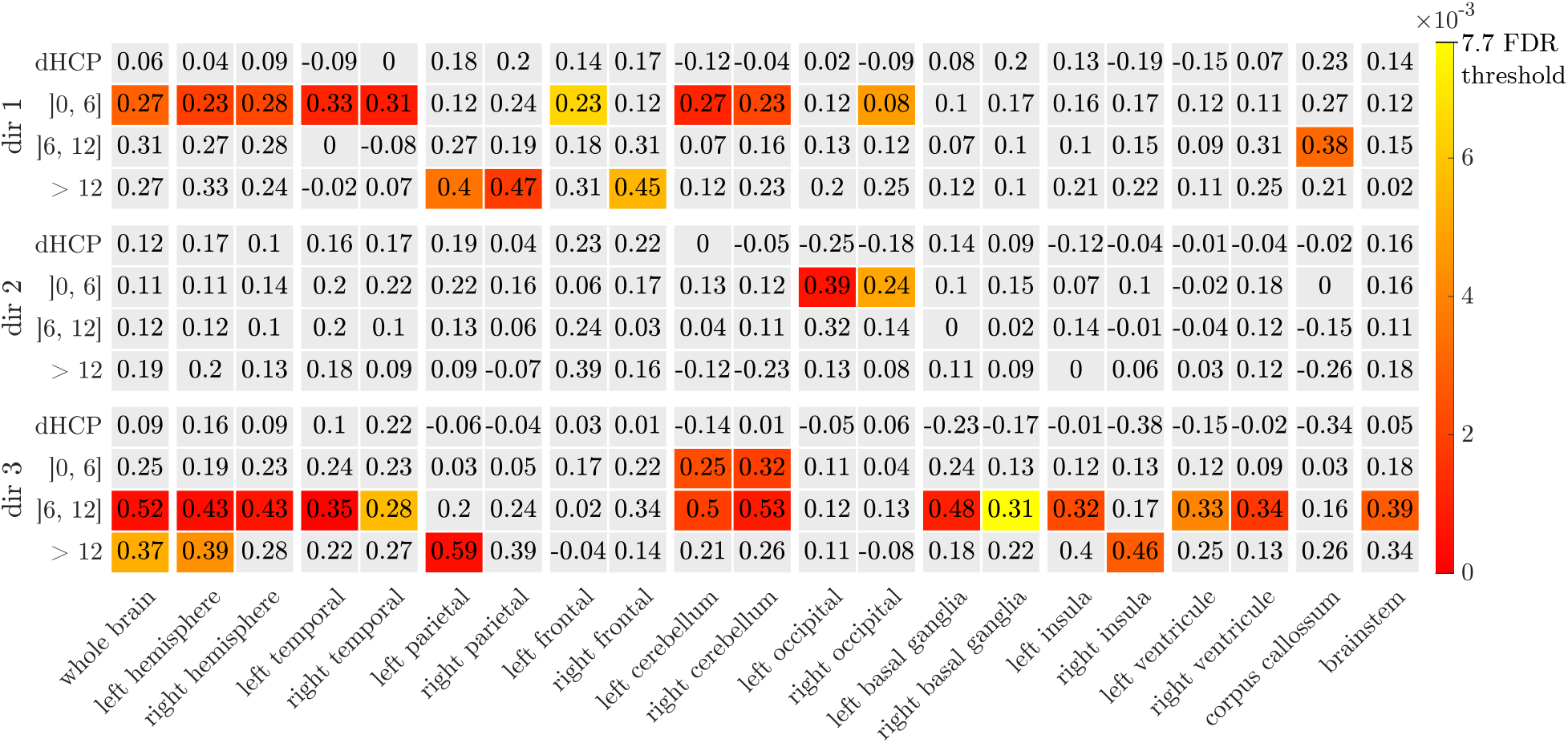
Male vs female comparison using Wilcoxon-Man-Whitney U-test and *H*_0_: the distrubution of the scaling factors of males and females are equal, Hi: the distributions of males and females are differents. In color: p-values for *H*_0_ rejection for FDR at level 5% (Benjamini and Hochberg method). Numericaly: the size of the effect *d* for each test.

A type 1 error, or false positive, occurs when *H*_0_ is incorrectly rejected. Since we are doing multiple comparisons, rejecting *H*_0_ based on the risk of type 1 error α = 5%, may lead in our case to an expected number of false positives superior to 12. Instead of using α, we therefore adopted the false discovery rate (FDR) that controls the proportion of false positives among the tests where *H*_0_ has been rejected. Therefore, we stated the acceptance or rejection of *H*_0_ based on a FDR at level 5%. This has been done using Benjamini and Hochberg procedure from [47] and corresponds to reject *H*_0_ when the p-value is less than 0.0077 (Figure 8). FDR has been preferred to family-wise error rate (FWER), that controls the risk of at least 1 false positive among the whole family of tests, because of the over-conservatism of this last type of procedure leading to poor test power (probability of correctly rejecting *H*_0_). Additionally, we calculated, for each test, the effect size *d* following: 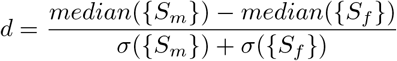 (Figure 8), where {*S_m_*} (resp. {*S_f_*}) is the set of scaling factor of males (resp. females) used for the test. We preferred the use of median instead of mean due to the fact that we do not know the distribution of the data a priori and we performed ranksum type tests.

For all the tests that lead to a rejection of the null hypothesis, scaling factors were higher for males both in terms of means and medians. Tests show that scaling factors of males seem higher in the second age class (0-6), brainwise and mainly in temporal and cerebellum areas along the direction 1. This is also notable in the same regions between 6 and 12 years, this time along direction 3. For the older class (12-19), this phenomenon essentially appears brainwise along the direction 3 and in the parietal lobes along direction 1.

### 4.4 Influence of the common reference

To evaluate the influence of the common reference image, the whole process described previously is reproduced using six different reference images. Those are atlases for different time-points *t*_1_,…, *t*_6_ based on the previously depicted population. Atlas for time *ti* is created using subjects with ages close to *t_i_* weighted according to their temporal distance to *t_i_* using kernel regression. Time-points are chosen such that five of them cover the period in which the majority of the brain expansion occurs, the last is positioned later, in a stabilized area (Figure 9).

**Fig 9.**
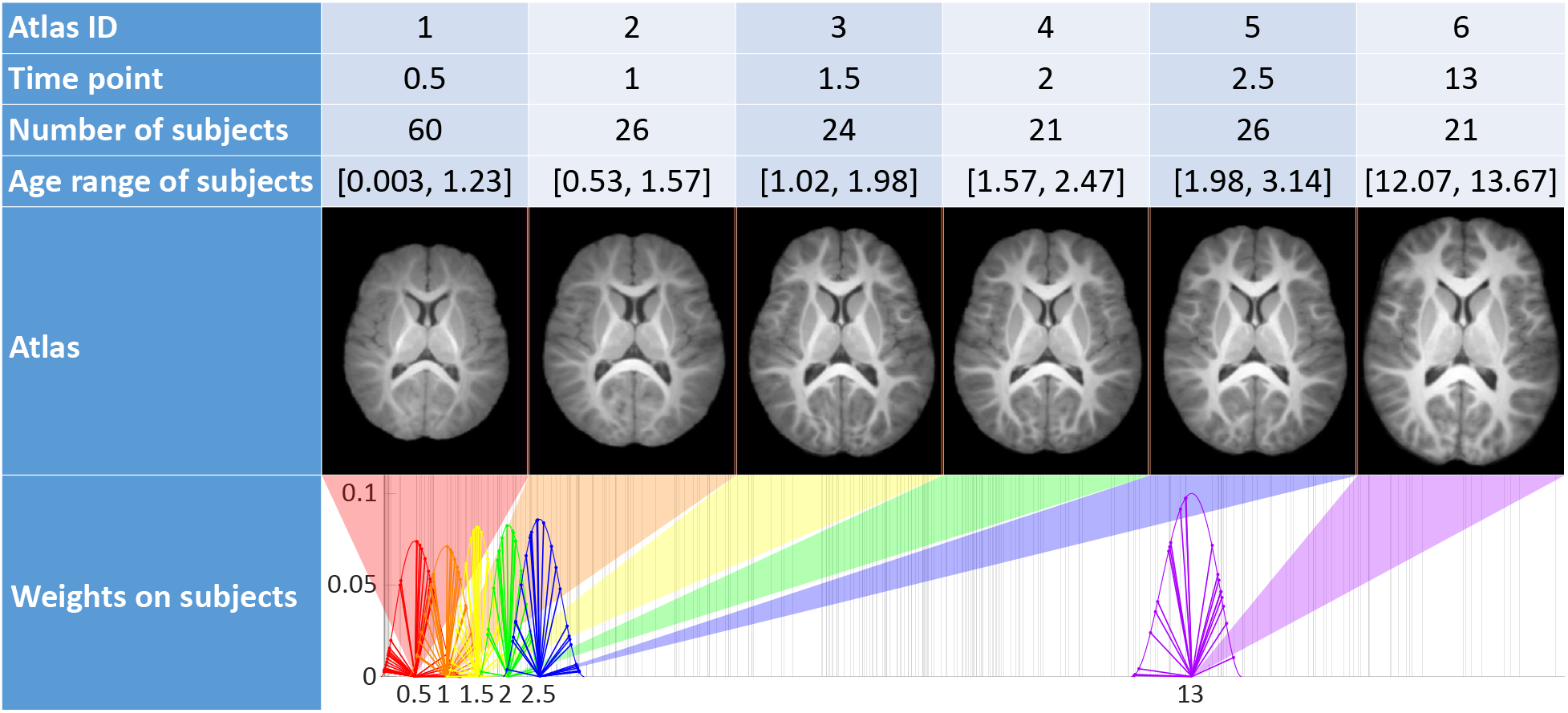
Characteristics of the 6 atlases used as reference image (time is displayed in years).

The method developed in section 3.2 is used for each of these reference images, on which directions of scaling for the anisotropic similarity registration have been established the same way. A scaling factor *s*(*i, j, d, k*) is thus computed for each ROI *j* of each subject i based on each reference image *k* along each chosen direction *d*. To quantify the influence of the reference image on absolute scaling factors, the results, using the six reference images previously depicted, are compared through two approaches:

1. A pairwise study to evaluate whether or not reference atlases closer to each other in age are more likely to generate closer results.
2. A study of the standard deviation among results for all reference atlases to evaluate how far they are from the average results.

#### 4.4.1 Study of pairwise distances between scaling factors by reference images in each direction for each ROI

Our aim is to determine whether or not reference images closer to each other (atlases at shorter temporal distances) are more likely to generate less important absolute differences between their results. We therefore to compute the absolute difference of the resulting scaling factors between each pairwise combinations of reference images. Then, those distances are normalized by the average of corresponding scaling factors between the two atlases such that it can be seen as a percentage of it (relative distance). The relative distance between scaling factors from reference atlases *k* and *l* is then computed as:

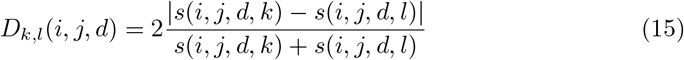

After examination of all the pairwise combinations, the temporal distance between the reference images does not seem to have an impact on the distance of the scaling ratios associated to each other (figure 10). The highest median of relative distance happens to be between atlases 2 and 5 for right basal ganglia, but does go above reach 8% of difference.

**Fig 10.**
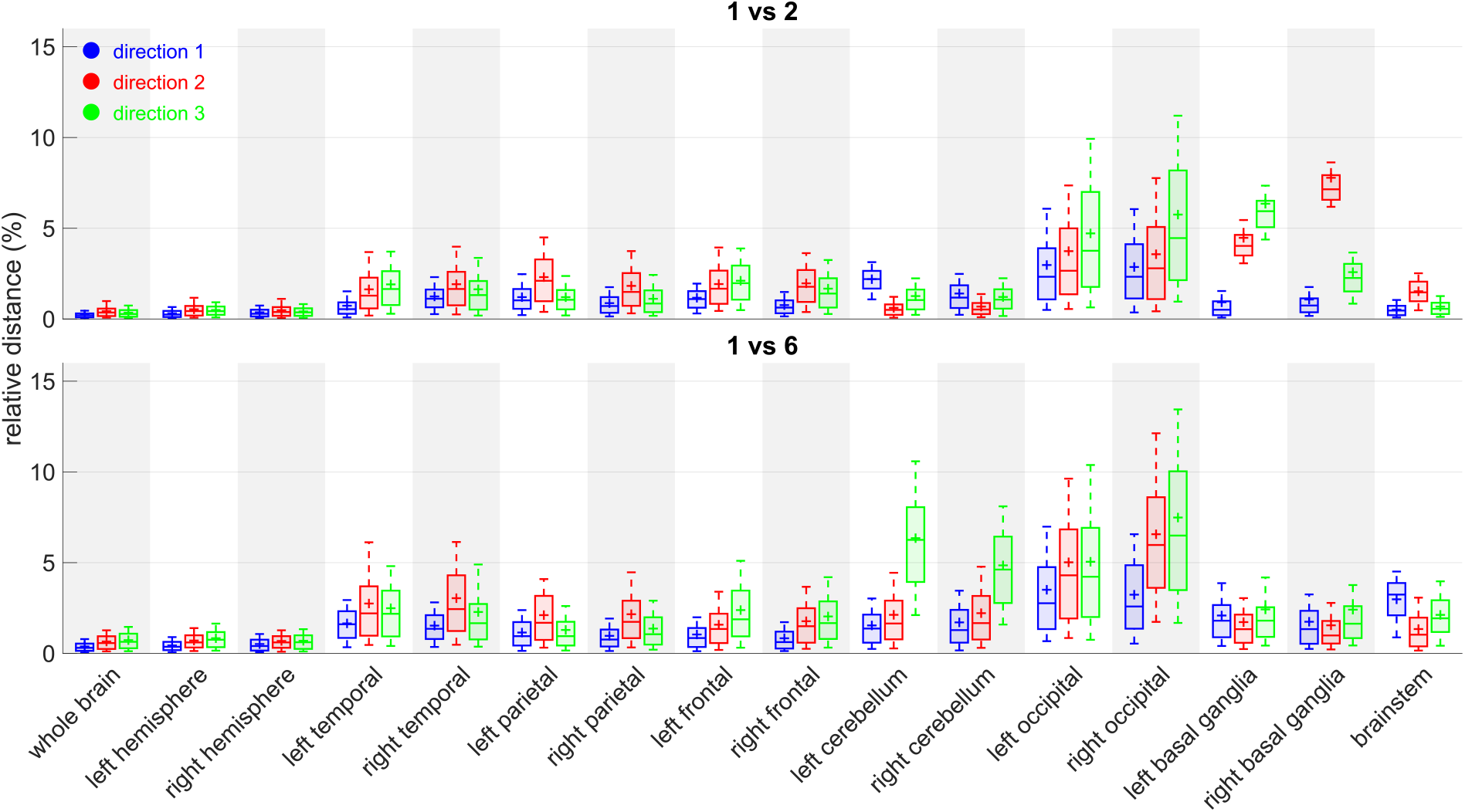
Relative distances between reference atlas 1 and 2 (top), 1 and 6 (bottom). Boxplots among subjects for each ROI *j*, each direction *d*: *boxplot*(*D_k,l_*(., *j, d*)).

#### 4.4.2 Study of the standard deviation among reference images in each direction for each ROI

This method gives an average measure of the distances between the results for each atlas and the average results. Those distances are normalized by the average of corresponding scaling factors of all the atlases. The relative standard deviation between scaling from all reference atlases is then computed as:

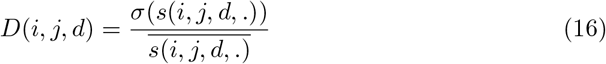

The graphs (figure 11) suggest that the method, when applied to large regions such as whole brain and hemispheres, is really robust to reference image change. Occipital lobes and cerebellum however seem to be more vulnerable areas. Those two regions share a common border and we believe that the segmentation process is a crucial step in that case. The cerebellum position indeed varies quickly in early stages of life and our decision to use segmentations based on neonates can be a bit inadequate for this area in particular. We also think that the way we chose to define the constrained directions of scaling (especially those using purely geometrical considerations through PCA on voxel coordinates) may not be the best suited for robustness in those areas. More anatomical features could lead to even smaller influence of the reference image.

**Fig 11.**
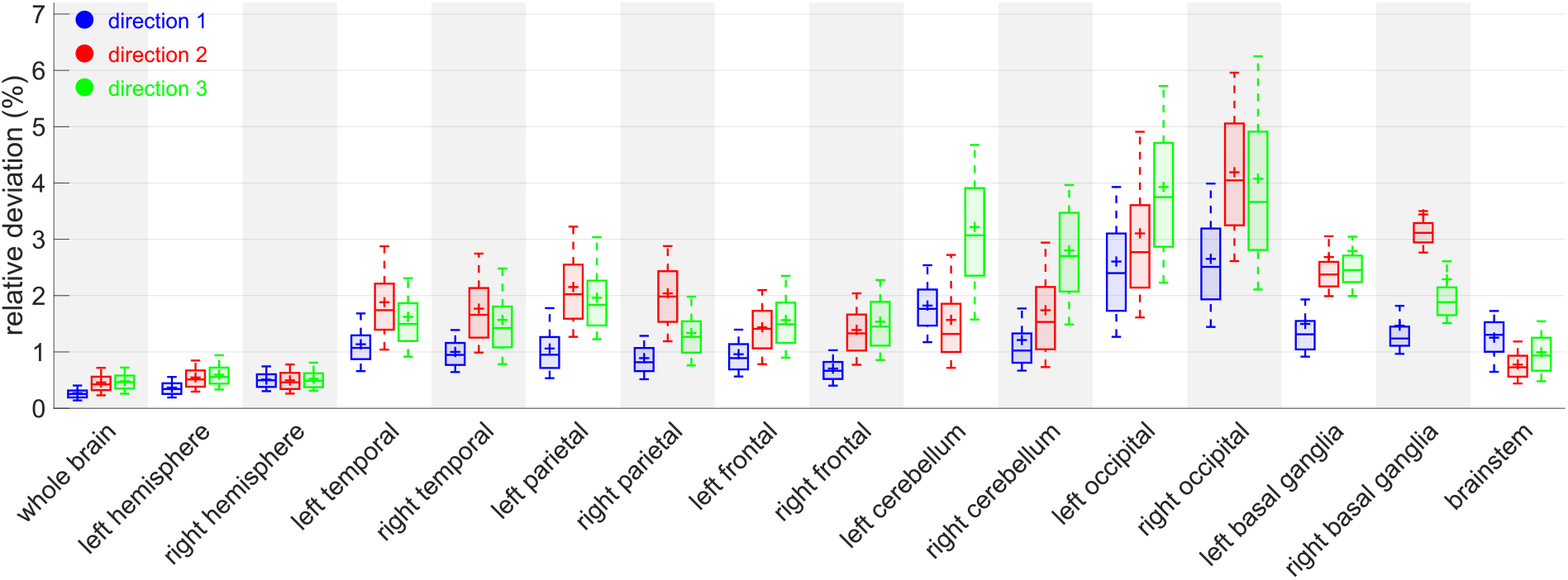
Relative standard deviation between reference atlases. Boxplots among subjects for each ROI *j*, each direction *d*: *boxplot*(*D*(., *j, d*)).

## 5 Discussion

The choice of the scaling directions, characterized by the matrix *U*, is crucial. If our decision to pick a vector orthogonal to the mid-sagittal plane seems meaningful due to symmetry reasons, the selection of the other two could be more debatable since they are based on purely geometric features ignoring iconic or anatomical considerations. A more anatomically-oriented approach could be to ask a medical expert to point, on the reference image, the anterior commissure - posterior commissure (AC-PC) line. This has been well adopted as a standard by the clinical neuroimaging community even though it is mostly a convention for visualization and at the cost of introducing a human interaction or a preprocessing step. There is no absolute good choice though and this choice depends on the purpose of the study. It is also possible to define specific directions for each ROI that could bring additional information for further studies. This method is therefore very flexible in the choice of the scaling directions and the ROIs, yet it has shown oneself robust concerning the choice of the common reference image.

We focused on the expansion of structures of a database of healthy subjects but we can also imagine using this method for patients. Intra-individual surveys are also possible, for subjects that had multiple scans through time, to monitor the evolution of a brain sub-region or any part of the body and infer the way it is going to expand.

Finally, although it does not call into question the method itself, there is room for improvements in the way we segmented the ROIs. The main difficulty is to find a method that is reproducible while being adaptable to brains from subjects scanned across a wide range of ages, which induces a large variability in contrast and shape.

## Conclusion

We have presented a method that allows the extraction of regional and global scaling factors along arbitrary chosen orthogonal directions. This is done through linear registration using a 9 dof transformation, anisotropic similarity, which is an affine transformation with constrained directions of scaling.

The main methodological contribution of this paper concerns the resolution of the problem of finding the optimal anisotropic similarity that best matches two sets of paired points. This result has made possible the development of a block-matching registration algorithm based on this transformation.

Given this new type of registration, our second contribution was to map a database of subjects between 0 and 19 years old using anisotropic similarity onto a common reference image on which the constrained directions of scaling of our choosing have been fixed. Registrations have been performed brainwise and ROI wise (lobes, cerebellum, basal ganglias…). For genericity purpose, we chose this reference image to be an atlas made from the subjects. Based on symmetry and geometrical considerations, we defined the same constrained directions of scaling for all ROIs even though it is possible to choose different ones for each. As an output, we obtained for each subject, for each ROI, for each chosen direction a scaling, a scaling factor that we normalized such that it represents an expansion factor from birth.

Those scaling factors have been used to model the anisotropic development of the brain. After model selection, it has been determined that rational function with polynomials of degree 1 as numerator and denominator is the best suited among the tested candidates for that modeling. Curves representing scaling factors as a function of the age for each ROIs, each chosen direction, along with associated confidence intervals have then beeen computed on a combination of four databases.

Tests to determine the influence of gender in those scaling factors have been performed for different age classes. Finally, two experiments have been conducted to evaluate the influence of the aforementioned common reference image. The results have shown small relative differences depending on the choice of the reference image leading to the conclusion that the method is robust in that aspect.

## 6 Acknowledgments

## 6.1 Databases

### 6.1.1 ASLpedia

A retrospective ASL study on routine pediatric MRI performed at Rennes University Hospital Neuropediatric radiology Department.

### 6.1.2 C-MIND

Data Repository created by the C-MIND study of Normal Brain Development. This is a multisite, longitudinal study of typically developing children from ages newborn through young adulthood conducted by Cincinnati Children’s Hospital Medical Center and UCLA and supported by the National Institute of Child Health and Human Development (Contract HHSN275200900018C). A listing of the participating sites and a complete listing of the study investigators can be found at: https://research.cchmc.org/c-mind.

### 6.1.3 The Developing Human Connectome Project (dHCP)

Led by King’s College London, Imperial College London and Oxford University, aims to make major scientific progress by creating the first 4-dimensional connectome of early life. https://data.developingconnectome.org/

### 6.1.4 ALBERTs

See [36] and [37] for details about segmentations. Copyright Imperial College of Science, Technology and Medicine and Ioannis S. Gousias 2013. All rights reserved. http://brain-development.org/brain-atlases/neonatal-brain-atlas-albert/

## A Detailed calculations for optimizing anisotropic similarity between two sets of paired points

## A.1

Let *R* be a rotation matrix. Then, an associated unit quaternion *q* is defined such as if 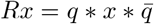. Thus:

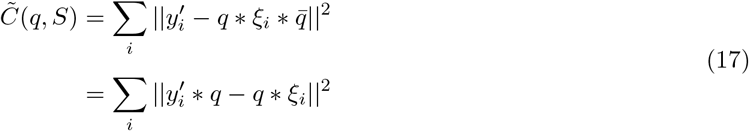

## A.2

If *p* is a vector, the associated quaternion is pure: *p*_1_ =0 which implies that *Q_p_* and *P_p_* are skew-symmetric. Yet 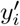 and *ξ_i_* are vectors, thus:

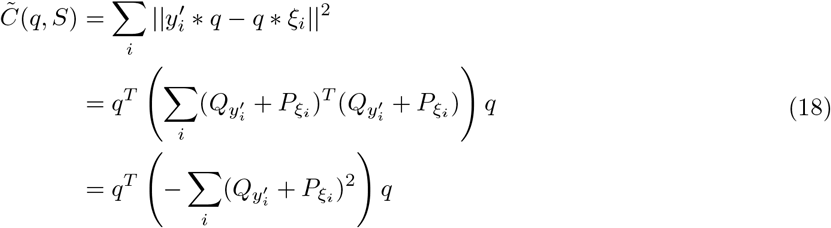

## A.3

If *p* is a vector, the associated quaternion is pure: *p*_1_ = 0 which implies that *Q_p_* and *P_p_* are skew-symmetric and 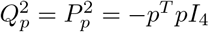.

Yet 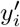 and *ξ_i_* are vectors, thus:

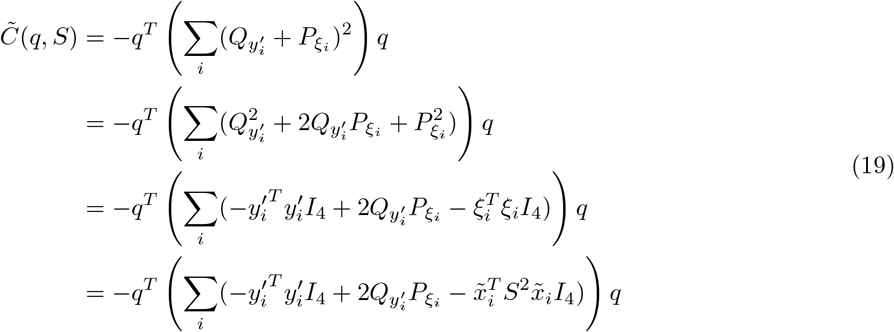

Thus:

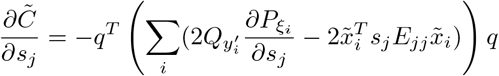

*E_jj_* being the matrix with a 1 at the intersection of the *j*^*th*^ row and the *j^th^* column and 0 elsewhere.

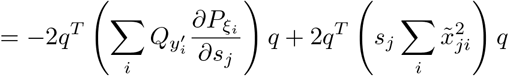

yet 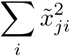 scalar and *q^T^q* = 1

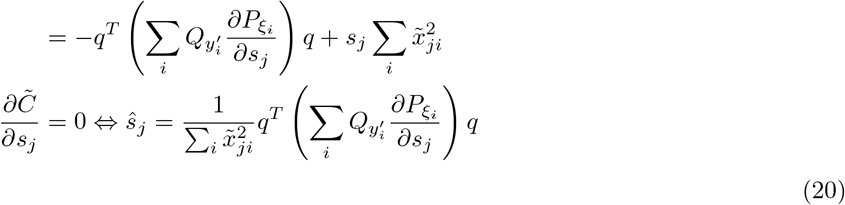

## A.4

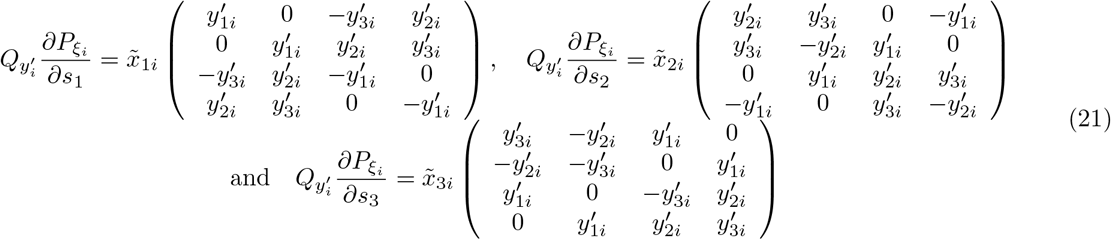

## Footnotes

1 Anima: github.com/Inria-Visages/Anima-Public/

2 Anima-Script: github.com/Inria-Visages/Anima-Scripts-Public/

3 Anima: github.com/Inria-Visages/Anima-Public/

